# Indirect and Direct Cannabinoid Agonists Differentially Affect Mesolimbic Dopamine Release and Related Behaviors

**DOI:** 10.1101/2020.06.14.150755

**Authors:** Kevin M. Honeywell, Timothy G. Freels, Megan A. McWain, Abigail S. Chaffin, Hunter G. Nolen, Helen J. Sable, Deranda B. Lester

## Abstract

The cannabinoid system is being researched as a potential pharmaceutical target for a multitude of disorders. The present study examined the effect of indirect and direct cannabinoid agonists on mesolimbic dopamine release and related behaviors in C57BL/6J (B6) mice. The indirect cannabinoid agonist *N*-arachidonoyl serotonin (AA-5-HT) indirectly agonizes the cannabinoid system by preventing the metabolism of endocannabinoids through fatty acid amide hydrolase (FAAH) inhibition while also inhibiting transient receptor potential vanilloid type 1 (TRPV1) channels. Effects of AA-5-HT were compared with the direct cannabinoid receptor type 1 (CB1R) agonist arachidonoyl-2’-chloroethylamide (ACEA). In Experiment 1, mice were pretreated with 7 daily injections of AA-5-HT, ACEA, or vehicle prior to assessments of locomotor activity using open field (OF) testing and phasic dopamine release using *in vivo* fixed potential amperometry. Chronic exposure to AA-5-HT did not alter locomotor activity or mesolimbic dopamine functioning. Chronic exposure to ACEA did not alter locomotor activity but did decrease phasic dopamine release while increasing the dopaminergic response to cocaine. In Experiment 2, mice underwent AA-5-HT, ACEA, or vehicle conditioned place preference (CPP) then saccharin preference testing, a measure commonly associated with anhedonia. Mice did not develop a CPP or aversion for AA-5-HT or ACEA, and repeated exposure to AA-5-HT or ACEA did not alter saccharin preference. Altogether, the findings suggest that neither of these drugs induce behaviors that are classically associated with abuse liability in mice; however, direct CB_1_R agonism may play more of a role in mediating mesolimbic dopamine functioning than indirect cannabinoid agonism.

---

The cannabinoid system is a promising target for pharmaceuticals treating a myriad of disorders including anxiety, depression, and addiction (Batista et al., 2014; Bortolato et al., 2007; Chen et al., 1990; Cohen et al., 2002; de Mello Schier et al., 2014; Economidou et al., 2006; Fattore et al., 2005; Fogaça, et al., 2012; Freels et al., 2020; Gobbi et al., 2005; Hakimizadeh et al., 2012; Li et al., 2009; Patel & Hillard, 2006; Rey et al., 2012; Xi et al., 2008; Zaitone et al., 2012). The neural impacts of drugs that act on the cannabinergic system are complex due to the abundance of cannabinoid receptors in the brain and the widespread interaction of endocannabinoids with other neurotransmitter systems. An improved understanding of the interactions between cannabinoids and other neurotransmitters should translate into more effective clinical uses for cannabinergic drugs. The present study examined the effect of indirect and direct cannabinoid agonists on mesolimbic dopamine release and related behaviors in mice.

The mesolimbic dopamine pathway consists of dopamine cell bodies in the ventral tegmental area (VTA) that project to limbic brain regions, predominately the nucleus accumbens (NAc). In freely moving rats, dopamine neurons fire tonically at ∼4 Hz and burst fire phasically at ∼20 Hz (Hyland et al., 2002). Tonic firing occurs without behaviorally relevant stimuli present and produces low concentrations of extracellular dopamine (Goto et al., 2007). Conversely, phasic dopamine firing occurs in relation to behaviorally significant external stimuli whose detection is crucial for learning, motivation, reward, and attention-shifting (Grace et al., 2007; Montague et al., 2004; Schultz et al., 1993). The mesolimbic dopamine system is often utilized to explore the addictive potential of drugs as most drugs of abuse and associated external cues increase phasic dopamine release in the NAc (Berridge & Robinson, 1998; Day et al., 2007; Everitt & Robinson, 2005).

The cannabinergic and dopaminergic systems are intertwined with receptor populations overlapping in multiple brain regions including the VTA and NAc (Han et al., 2017; Lupica & Riegel, 2005; Szabo et al., 2002; Winters et al., 2012). The cannabinergic system consists of two primary types of cannabinoid receptors, type 1 (CB_1_Rs) and type 2 (CB_2_Rs). Postsynaptic neuron depolarization triggers the release of endocannabinoids including *N*-arachidonoyl ethanolamide (AEA) that binds to CB_1_Rs and 2-arachidonoylglycerol (2-AG) that binds to CB_1/2_Rs, which both inhibit presynaptic neurotransmission before being metabolized by fatty acid amide hydrolase (FAAH) and monoacylglycerol lipase (MAGL), respectively (Batista et al., 2014; Howlett et al., 2002; Kano et al., 2009; Blankman & Cravatt, 2013; Dinh et al., 2002; Giang & Cravatt, 1997; Gulyas et al., 2004; Haj-Dahmane, 2018; Kaczocha et al., 2009; Ohno-Shosaku et al., 2001; Maccarrone, 2017). Select cannabinoids, including the endocannabinoid AEA, also act on the transient receptor potential vanilloid (TRPV) channels, which can be located pre- and post-synaptically to regulate neurotransmission in mesolimbic brain regions (De Petrocellis et al., 2001; Kaur & Gibson, 2009; Smart et al., 2000; Rosenbaum & Simon, 2007; Ross, 2003). Thus, cannabinergic drugs have multiple mechanisms by which to alter mesolimbic dopamine release. The present study examined the dopaminergic effects of an indirect CB_1_R agonist (*N*-arachidonoyl serotonin: AA-5-HT) and a direct CB_1_R agonist (arachidonoyl-2’-chloroethylamide: ACEA). AA-5-HT indirectly agonizes the cannabinoid system by preventing the metabolism of endocannabinoids through FAAH inhibition while also inhibiting TRPV type 1 (TRPV_1_) channels (Maione et al., 2007; Micale et al., 2009).

The influence of cannabinoid agonists on mesolimbic dopamine functioning is controversial in the literature with conflicting findings likely due to differences in techniques used to measure dopamine release and differences in drugs employed and respective doses to alter cannabinergic and dopaminergic signaling. Using microdialysis, AEA administration increased tonic dopamine levels in the NAc, an effect that was attenuated by a CB_1_R antagonist (Solinas et al., 2006). In this same study, an FAAH inhibitor increased AEA-induced tonic dopamine elevations in the NAc, and this effect was also attenuated by CB_1_R antagonism. Others have shown similar results with Δ^9^-tetrahydrocannabinol (THC) increasing tonic dopamine levels and CB_1_R antagonism blocking the effect (Tanda et al., 1997). Self-administration of a direct CB_1_R agonist also increased NAc tonic dopamine release (Fadda et al., 2006; Lecca et al., 2006). Most studies examining the influence of cannabinoids (CBs) on tonic dopamine levels agree that CB agonists increase tonic dopamine release by utilizing CB_1_Rs; however, the effects of CB agonists on phasic dopamine release are less clear. We have previously shown that the administration of either AA-5-HT or ACEA attenuated stimulation-evoked phasic dopamine release in the NAc as measured by fixed potential amperometry with the direct CB_1_R agonist ACEA having a more extensive effect than the indirect agonist AA-5-HT (Freels et al., 2020). A study utilizing fast scan cyclic voltammetry (FSCV) found that while CB_1_R agonism decreased stimulation-evoked dopamine release in the NAc the number of spontaneous phasic transients per minute was increased (Cheer et al., 2004). Drugs targeting the CB system can also alter the effects of other drugs that either indirectly or directly acts on the mesolimbic dopamine pathway. For example, a CB_1_R antagonist has been shown to attenuate cocaine’s effects on both basal and phasic dopamine release in the NAc (Cheer et al., 2007; Mereu et al., 2013), and Peters et al. (2021) found that direct CB_1_R antagonism decreased tonic and phasic dopamine release evoked by reward-associated cues while an FAAH inhibitor had no effect.

Additionally, CB agonists alter behaviors related to dopamine transmission. Previous research has shown that administration of a direct CB_1_R agonist either did not affect or reduced locomotion and either did not affect or reduced rearing (Freels et al., 2020; Cheer et al., 2004; Vlachou et al., 2008). CB manipulation also alters the behavioral effect of dopaminergic drugs. Pretreatment with a CB_1_R antagonist reduced cocaine-induced hyper locomotion while pretreatment with an FAAH inhibitor had the opposite effect (Mereu et al., 2013). Similarly, genetic deletion and pharmacological blockade of CB_1_Rs have been shown to reduce cocaine self-administration (Soria et al., 2005), and CB_1_R agonists have been shown to increase cocaine seeking (De Vries et al., 2001; Friedman et al., 2019). Such studies indicate a faciliatory role for CBs in cocaine reward. However, others have shown that CB_1_R agonists reduced cocaine-induced locomotion and cocaine self-administration (Fattore et al., 1999; Vlachou et al., 2003; Vlachou et al., 2008), suggesting that in some contexts CB agonism may reduce the reinforcing effects of rewarding stimuli. As such, in both human and rodent studies, chronic cannabis exposure has been linked to impaired motivation and anhedonia (see Blum et al., 2021).

The current study aimed to determine the effects of repeated exposure to the indirect CB agonist AA-5-HT and the direct CB_1_R agonist ACEA on mesolimbic dopamine release and related behaviors, specifically locomotor activity, conditioned place preference (CPP), and anhedonia. The study consisted of 2 experiments. In Experiment 1, mice were pretreated with 7 daily injections of AA-5-HT, ACEA, or vehicle (negative control) prior to assessments of locomotor activity using open field (OF) testing and dopamine release using *in vivo* fixed potential amperometry (*Figure 1A)*. In Experiment 2, mice underwent AA-5-HT, ACEA, or vehicle CPP for eight days (same dosing regimen as used in Experiment 1) followed by saccharin preference testing, a measure commonly associated with anhedonia (*Figure 1B)*. The current study is an extension of our previous study showing that acute administrations of AA-5-HT and ACEA decrease dopamine release (Freels et al., 2020). The chronic drug administration employed in the current study is more generalizable to human drug CB use. Furthermore, in the current study some mice received an administration of cocaine during the amperometric dopamine recordings, allowing us to address the aforementioned cross-sensitization effects between the cannabinergic and dopaminergic systems.

**Figure 1.**
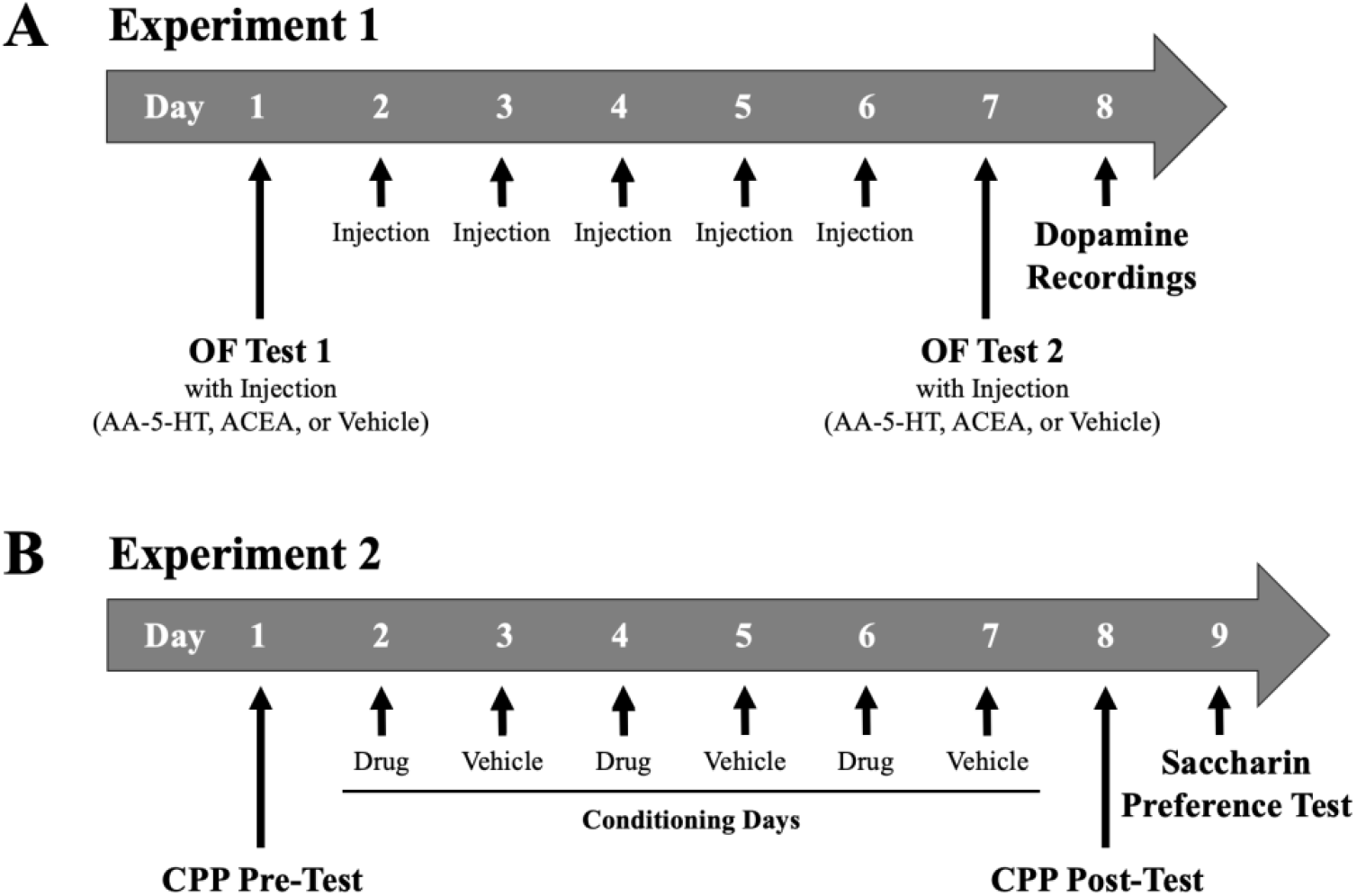
Experimental Timelines. *Note.* Experimental timelines are depicted. (A) In Experiment 1, mice were pretreated with 7 daily injections of AA-5-HT, ACEA, or vehicle (negative control) prior to assessments of locomotor activity using OF testing and dopamine release using *in vivo* fixed potential amperometry. (B) In Experiment 2, mice underwent AA-5-HT, ACEA, or vehicle CPP for eight days followed by saccharin preference testing, a measure commonly associated with anhedonia. OF = open field, AA-5-HT = arachidonoyl serotonin, ACEA = arachidonyl-2’-chloroethylamide, and CPP = conditioned place preference.

## Methods

All procedures have been approved by the Institutional Animal Care and Use Committee (IACUC) at the University of Memphis and were also aligned with those outlined in *The Public Health Service Policy on Humane Care and Use of Laboratory Animals* (National Institutes of Health, 2012) and the *Guidelines for the Care and Use of Mammals in Neuroscience and Behavioral Research* (National Research Council, 2003).

### Animals

Fifty-four, male C57BL/6J (B6) mice for Experiment 1 and 27, male B6 mice for Experiment 2 were obtained from Jackson Laboratory (Bar Harbor, ME) and were housed 2-5 per cage in 18.5 x 29.5 cm polycarbonate Generic State Microisolators with Sani-Chips bedding (P.J. Murphy Forest Products Montville, NJ). Food and water were available *ad libitum*. Mice were housed in a temperature-controlled room (21 ± 1° C) with a 12 h light: 12 h dark cycle. At the time of drug treatments and experiments, mice were 3 to 5 months old (adults).

### Experiment 1 (Locomotor Activity and Dopamine Release)

#### Chronic Drug Treatments

Chronic drug treatments consisted of one i.p. injection per day for seven days of either AA-5-HT (2.5 mg/kg, *n* = 19), ACEA (1.0 mg/kg, *n* = 15), or vehicle comprised of 10% dimethyl sulfoxide (DMSO) in saline (*n* = 20) all acquired from Sigma Aldrich (St. Louis, Missouri). On the eighth day, during dopamine recordings, mice received an i.p. injection of AA-5-HT (2.5 mg/kg), ACEA (1.0 m/kg), cocaine (10.0 mg/kg), or vehicle. The ten total groups encompassed AA-5-HT:AA-5-HT (*n* = 7), AA-5-HT:vehicle (*n* = 6), AA-5-HT:cocaine (*n* = 6), ACEA:ACEA (*n* = 5), ACEA:vehicle (*n* = 5), ACEA:cocaine (*n* = 5), vehicle:AA-5-HT (*n* = 6), vehicle:ACEA (*n* = 4), vehicle:vehicle (*n* = 4), and vehicle:cocaine (*n* = 6) where the first drug represents that given during behavioral testing and the second drug represents that given during dopamine recordings. To minimize outside anxiogenic variables, drug treatments were administered by the same person, at the same time (during the light portion of the light/dark cycle) each day.

#### OF Testing

Mice were tested in the OF chamber twice once on the first and once on the seventh day of the chronic drug treatment. On test day, each mouse was placed in a single holding cage inside a sound attenuated cabinet in the testing room for 45 min in order to habituate the mouse. At the beginning of each OF test, the mouse was placed in the center of the OF chamber. The OF apparatus was a HamiltonKinder SmartFrame™ (HamiltonKinder, Poway, CA) with a clear Plexiglas insert with dimensions of 24.13 cm x 45.72 cm, a 4 x 8 photo beam strip, and a 4 x 8 photo beam rearing attachment. During the session, software (MotorMonitor verison 4.14, HamiltonKinder, Poway, CA) tracked the time spent in the central area, rearing, and total distance travelled. The central area specified in the software’s zone map function was a 9 cm x 10 cm space positioned 4.5 cm from the left and right walls and 15 cm from the front and back walls of the chamber. After a 20 min baseline movement assessment, the mouse received an i.p. injection of the assigned chronic treatment (AA-5-HT, ACEA, or vehicle) and testing continued for 90 min post-injection. At the end of the test, each mouse was returned to its home cage in the mouse colony. The OF chamber was then cleaned with 10% isopropyl alcohol and allowed to dry after each trial.

#### Dopamine Recordings

On the eighth day of the chronic drug treatments, all mice underwent stereotaxic surgery for the measurement of dopamine transmission using *in vivo* fixed potential amperometry. Mice were permanently anaesthetized with urethane (1.5 g/kg, i.p.). The dose of which was split into two i.p. injections separated by 10 min. Mouse anesthesia was assessed through eye blink, mild tail pinch, and mild foot pinch reflexes. Mice were mounted into a stereotaxic frame (David Kopf Instruments, Tujunga, CA), and their body temperature was kept at approximately 37°C. The stimulating electrode (SNE-100; Rhodes Medical Co., Summerland, CA) was placed into the left VTA (coordinates in mm from bregma: AP −3.3, ML +0.3, and DV −4.0 from dura) (Paxinos & Franklin, 2001). The Ag/AgCl reference and stainless-steel auxiliary electrode combination was placed on contralateral cortical tissue −2.0 mm to bregma, and the carbon fiber recording electrode (500 um length x 7 um o.d.; Union Carbide, North Seadrift, TX) was positioned in the left NAc (coordinates in mm from bregma: AP +1.5, ML +1.0, and DV −4.0 from dura) (Paxinos & Franklin, 2001). A series of cathodal current pulses was delivered to the stimulating electrode via an optical stimulus isolator and programmable pulse generator (Iso-Flex/Master 8; AMPI, Jerusalem Israel), which stimulated dopamine release. A fixed +0.8 V potential was continuously applied to the recording electrode, which oxidized the dopamine. The change in current due to the oxidation of dopamine was monitored by the electrometer (ED401 e-corder and EA162 Picostat, eDAQ Inc., Colorado Springs, CO) sampling 10k times per sec and filtered at 50 Hz.

Stimulation parameters varied depending on the aspect of dopamine transmission being measured. Initially, while establishing a baseline response, the stimulation protocol consisted of 20 monophasic 0.5 ms duration pulses (800 µAmps) at 50 Hz. Dopamine autoreceptor (DAR) sensitivity was assessed by applying a pair of test stimuli (T1 and T2, each 10 pulses at 50 Hz with 10 s between T1 and T2) to the VTA every 30 s (Fielding et al., 2013; Holloway et al., 2018; Mittleman et al., 2011). Seven sets of conditioning pulses (0, 1, 5, 10, 20, 40, and 80; 0.5 ms pulse duration at 15 Hz) were delivered prior to T2 such to leave 0.3 s between the end of the conditioning pulse train and initiation of T2. DAR-mediated inhibition of evoked dopamine efflux was expressed in terms of the change in the amplitude of T2 with respect to T1 for each set of conditioning pulses where low-to-high DAR sensitivity was represented as low-to-high percent inhibition of evoked dopamine efflux (i.e. high sensitivity results in lower amplitude of T2 relative to T1).

Upon completion of the DAR sensitivity test, stimulation parameters were reset to 20 pulses at 50 Hz every 30 sec. Following 5 min of baseline dopamine efflux recording, each mouse was given a drug challenge via an i.p. injection of AA-5-HT (2.5 mg/kg), ACEA (1.0 mg/kg), cocaine (10 mg/kg), or vehicle. Dopamine recordings continued for 90 min post drug challenge for all groups. After the recordings were complete, direct anodic current of 100 µAmps was applied to the stimulating electrode for 10 s to create an iron deposit, which marked the electrode’s position.

Mice were euthanized via an intracardial injection of urethane (0.345 g/mL). Brains were removed and stored in 30% sucrose / 10% formalin solution with 0.1% potassium ferricyanide. Coronal sections of each brain were sliced at −30°C using a cryostat, and electrode placements were identified using a light microscope and marked on coronal diagrams (Paxinos & Franklin, 2001). Following the experiment, *in vitro* electrode calibration occurred by recording in solutions of dopamine (0.2 µM – 1.2 µM) via a flow injection system (Dugast et al., 1994; Prater et al., 2018), which allowed for the conversion of current measurements, to dopamine concentrations.

### Experiment 2 (CPP and Saccharin Preference Test)

#### AA-5-HT or ACEA CPP

Experiment 2 aimed to determine whether mice would establish a CPP (or a conditioned place aversion; CPA) for the side paired with either the indirect or direct cannabinoid agonist (AA-5-HT or ACEA, respectively). Prior to each session, each mouse was placed in a single holding cage inside a sound attenuated cabinet in the testing room for 45 min to minimize outside anxiogenic variables. On the first day (habituation), the mouse was placed into the HamiltonKinder SmartFrame™ (HamiltonKinder, Poway, CA) equipped with an insert to divide the OF into two halves measuring 21.84 cm x 22.23 cm each. There was a center doorway between the two chambers measuring 6.99 cm (width) x 8.89 cm (height). The walls of the chamber were covered with either vertical or horizontal black and white bars each measuring

2.54 cm thick. The vertical bars side was paired with Sani-chips bedding (P.J. Murphy Forest Products, Montville, NJ), and the horizontal bars side was paired with So Phresh Natural Softwood bedding (Petco Animals Supplies, San Diego, CA). A black grating covered the beddings. Mice were allowed to access both chambers for the 30 min test period. On conditioning days 2-7, a black, opaque insert was placed over the doorway to restrict movement during the 30 min sessions to only one side. On conditioning days 2, 4, and 6, each mouse was given an i.p. injection of either AA-5-HT (2.5 mg/kg, *n* = 9), ACEA (1.0 mg/kg, *n* = 9), or vehicle (*n* = 9), while, on conditioning days 3, 5, and 7, all mice received vehicle. The drug-paired side was counterbalanced for all treatment groups. On the eighth day (test day), the mouse was again placed into the chamber with the door insert removed for 30 min. At the end of each session, each mouse was returned to its home cage in the colony room, and the chamber was cleaned with 10% isopropyl alcohol and allowed to dry.

#### Two-Bottle Saccharin Choice Test

Additionally, for the second experiment, one day following CPP testing, mice underwent a two-bottle choice test for saccharin or water preference. A reduction in the preference ratio for saccharin during this test is indicative of anhedonia (Sheffield & Roby, 1950). At lights on, the mice were weighed, and the water bottles were removed from their home cages. At lights off (12 hours later), the mice were injected with AA-5-HT (2.5 mg/kg, *n* = 9), ACEA (1.0 mg/kg, *n* = 9), or vehicle (*n* = 9) and placed individually into separate cages each containing one bottle of 0.1% saccharin and one bottle of tap water (both weighed beforehand). After the 2 h test, mice were returned to their home cage with the home cage water bottle replaced. The preference test bottles were then weighed again to determine how much of each had been consumed. To eliminate novelty effects, only the second to sixth day were used to examine group differences.

### Data Analyses

#### Data Analyses for Experiment 1 (Locomotor Activity and Dopamine Release)

For the first experiment, OF data was analyzed using three-way ANOVAs for day x block x treatment effects, which includes distance travelled, rears, and percent time in center. Significant treatment differences indicated by *p* < .05 were further explored using Tukey’s HSD post-hoc tests or Games-Howell post-hoc tests when appropriate. Electrochemical data were broken down into DAR sensitivity, baseline dopamine recordings, and drug challenge recordings. Dopamine oxidation current recordings were used to quantify VTA stimulation induced dopamine release in the NAc by extracting data points occurring between 0.25 s pre- and 10 s post-stimulation at the desired time. DAR-mediated inhibition of evoked dopamine release was expressed in terms of the percent change between test stimulations (T2/T1×100) for each set of pre-pulses (Fielding et al., 2013; Holloway et al., 2018; Mittleman et al., 2011). One vehicle-vehicle mouse was removed from this analysis due to technical issues. A two-way mixed ANOVA was used to assess the impact of chronic treatment (AA-5-HT, ACEA, or vehicle) and number of pre-pulses on percent inhibition. From baseline (pre-drug challenge) recordings, dopamine release was quantified as the magnitude of the response, and dopamine synaptic half-life, an indication of dopamine transporter (DAT) functioning, was quantified as the time for 50% decrease from the maximum evoked increase to the pre-stimulus level (Holloway et al., 2018; Mittleman et al., 2011). One-way ANOVAs, with Tukey’s HSD post-hoc tests as needed, were used to determine the effect of chronic, drug treatment on baseline dopamine release and half-life. During dopamine recordings, mice were administered a drug challenge of AA-5-HT, ACEA, cocaine, or vehicle. Following the drug challenge, data was extracted at 10 min intervals. Change in dopamine release and half-life was expressed as percent change relative to baseline release and half-life (pre-drug = 100%). For each drug challenge, a two-way mixed ANOVA was used to assess the impact of chronic, drug treatment and time post injection on percent change in dopamine release and half-life. A one-way ANOVA with Tukey post hoc analysis was used when appropriate to test the effects of chronic, drug treatment on percent change in dopamine release or half-life at each time point. Significant differences indicated by *p* < .05 were further explored using Games-Howell post-hoc tests when appropriate.

#### Data Analyses for Experiment 2 (CPP and Saccharin Preference)

For the second experiment, the two sets of behavioral data were analyzed separately. For place preference, potential chamber bias was first examined using a one sample *t*-test comparing the time spent on what was to become the drug-paired side to 900 s (i.e., exactly half of the habituation session time). Next, time spent on the drug-paired side on day one was subtracted from the time spent on the drug-paired side on day eight, and the difference was analyzed by a one-way ANOVA to determine if a CPP or CPA was present. Likewise, the number of entries into the drug-paired side on day one was subtracted from the number of entries into the drug-paired side on day eight, and the difference was analyzed by a one-way ANOVA. Locomotor activity was also examined by looking at the total distance travelled (cm) on each of the three drug-paired days and analyzing these data with a two-way ANOVA. For the two-bottle choice test, saccharin consumed in g/kg body weight and saccharin preference (i.e., percent of total liquid consumed) were analyzed separately by two-way ANOVA.

## Results

### Experiment 1 (Locomotor Activity and Dopamine Release)

#### OF Testing

Locomotor activities assessed in the OF included distance travelled (cm), rearing, and percent time in center. Baseline locomotor activity was assessed in two 10 min blocks per day. As can be seen in Figure 2A and Figure 2B, there was not a day x block x chronic, drug treatment interaction effect, day x chronic drug treatment interaction effect, block x chronic, drug treatment interaction effect, or a main effect of chronic, drug treatment on distance travelled (all *p*’s > .05). As can be seen in Figure 2C and Figure 2D, there was also no interaction effects or main effect of chronic, drug treatment on baseline rearing (all *p*’s > .05). As can be seen in Figure 2E and Figure 2F, there was also no interaction effects or main effect of chronic, drug treatment on baseline percent time in center (all *p*’s > .05).

**Figure 2.**
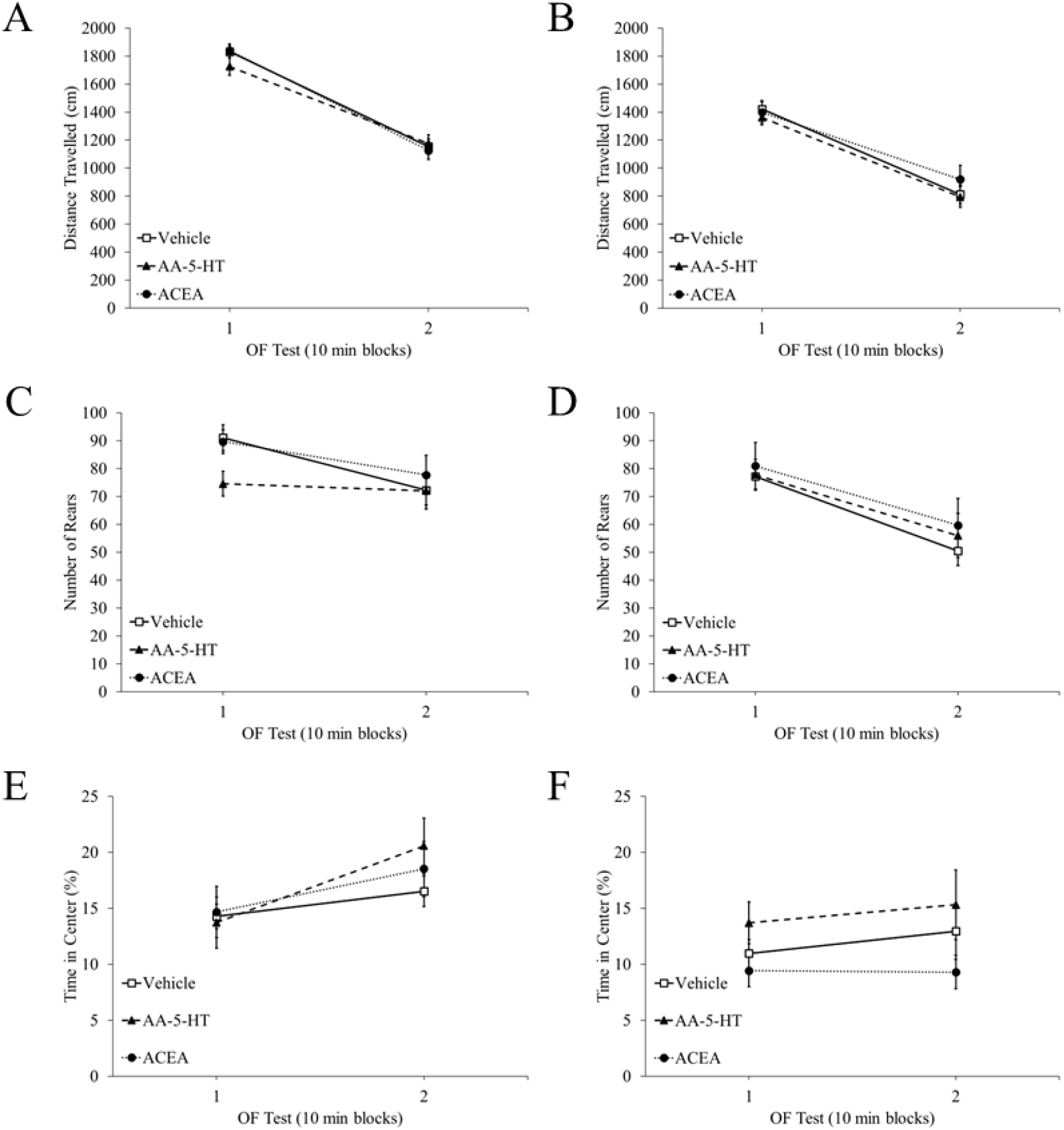
Baseline Locomotor Activities. *Note.* Baseline differences in locomotor activities in the OF were averaged into 10 min blocks and did not differ on (A) distance travelled on OF test 1, (B) distance travelled on the OF test 2, (C) number of rears on OF test 1, (D) number of rears on OF test 2, (E) time spent in the center on OF test 1, or (F) time spent in the center on OF test 2. OF = open field, AA-5-HT = arachidonoyl serotonin, and ACEA = arachidonyl-2’-chloroethylamide.

During the drug challenge in the OF on the first and seventh day, locomotor activities were recorded in 10 min blocks over the 90 min. As can be seen in Figure 3A and Figure 3B, there was not a day x block x chronic, drug treatment interaction effect, day x chronic, drug treatment interaction effect, block x chronic, drug treatment interaction effect, or main effect of chronic, drug treatment on distance travelled (all *p*’s > .05). As can be seen in Figure 3C and Figure 3D, there was a main effect of chronic, drug treatment on rears, *F*(2,51) = 4.53, *p* = .02, η_p_ ^2^ = .15. Games-Howell post-hoc test revealed that ACEA pretreated mice (*M* = 5.50, *SE* = 0.04) reared significantly less than vehicle pretreated mice (*M* = 11.51, *SE* = 0.05), *p* = .01. However, there were no interactions of chronic, drug treatments with block or day (all *p*’s > .05). As can be seen in Figure 3E and Figure 3F, there was also no interaction effects or main effect of chronic, drug treatment on percent time in the center (all *p*’s > .05).

**Figure 3.**
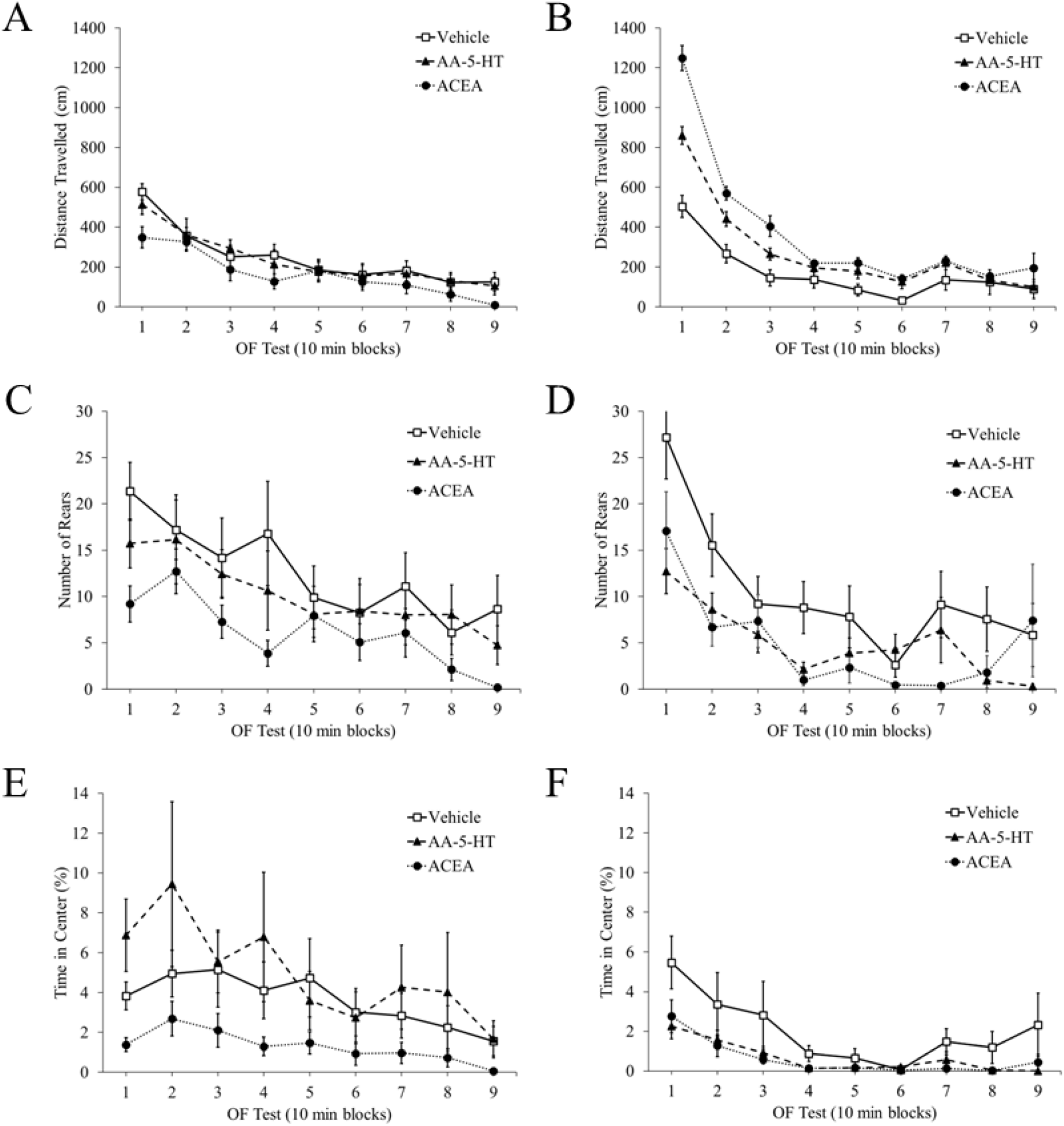
Drug Challenge Locomotor Activities. *Note.* Differences following a drug challenge in locomotor activities in the OF were averaged into 10 min blocks and did not differ on (A) distance travelled on OF test 1, (B) distance travelled on OF test 2, (C) number of rears on OF test 1, (D) number of rears on OF test 2 even though ACEA significantly reduced overall rears, (E) time spent in the center on OF test 1, or (F) time spent in the center on OF test 2. Though, there was a main effect chronic, drug treatment on rears. OF = open field, AA-5-HT = arachidonoyl serotonin, and ACEA = arachidonyl-2’-chloroethylamide.

#### Dopamine Recordings Stereotaxic Placement of Electrodes

As can be seen in Figure 4A, following amperometry experiments, the placement of stimulating electrodes (*n* = 54) and recording electrodes (*n* = 54) were determined by examining lesioned regions in sectioned mouse brains. The positions of stimulating electrodes were localized within the anatomical region of the VTA spanning −3.08 to −3.52 mm AP from bregma and −4.0 to −5.0 mm DV from dura, and the positions for the recording electrodes were localized within the anatomical region of the NAc spanning +1.54 to + 1.34 mm AP from bregma and −4.0 to −5.0 mm DV from dura.

**Figure 4.**
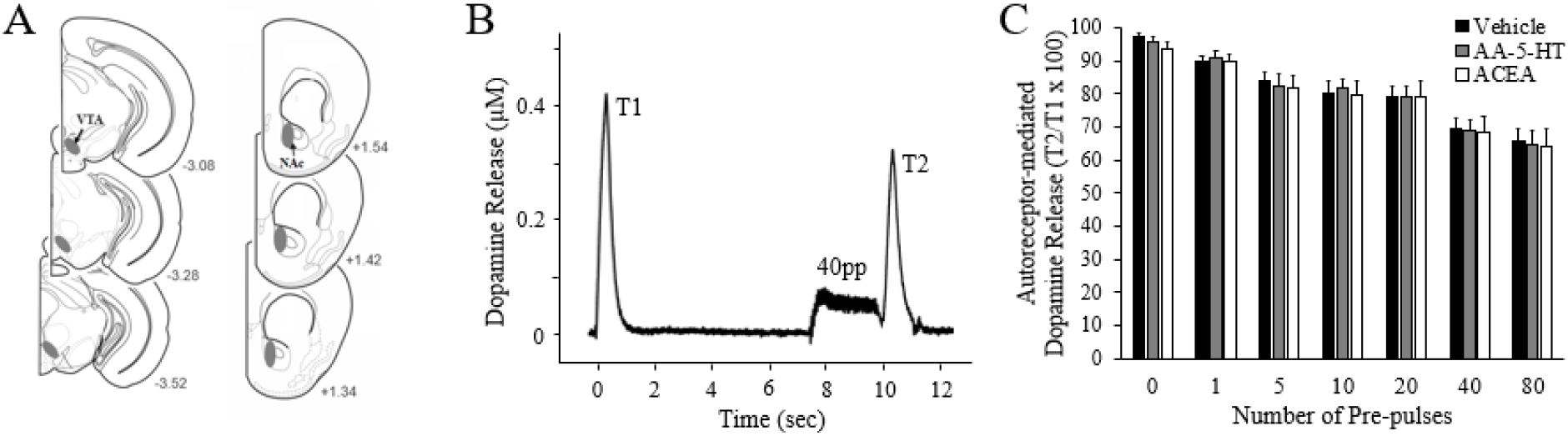
Electrode Placement and Dopamine Release Inhibition. *Note.* (A) Representative coronal sections of the mouse brain (adapted from the atlas of Paxinos & Franklin, 2001), with gray-shaded areas indicating the placements of stimulating electrodes in the ventral tegmental area (VTA) and amperometric recording electrodes in the nucleus accumbens (NAc). Numbers correspond to mm from bregma. (B) A representative amperometric recording during the autoreceptor functioning test and (C) autoreceptor-mediated dopamine release displayed as means ± SEMs following pre-pulses. Chronic pretreatment with either AA-5-HT (arachidonoyl serotonin) or ACEA (arachidonyl-2’-chloroethylamide) did not alter dopamine autoreceptor functioning in the NAc.

##### DAR Sensitivity

DAR sensitivity was analyzed across chronic, drug treatment conditions as previously described (Fielding et al., 2013; Holloway et al., 2018; Mittleman et al., 2011). With a representative recording seen in Figure 4B and means with standard error in Figure 4C, there was not a main effect of chronic, drug treatment on DAR sensitivity, *F*(2,50) = 0.08, *p* = .92, η_p_^2^ = .00. According to Mauchly’s test, the assumption of sphericity had been violated for number of pre-pulses, χ^2^(20) = 180.42, *p* < .001, and, therefore, degrees of freedom were corrected using Greenhouse-Geisser estimates of sphericity (ε = 0.39). As expected, there was a main effect of number of pre-pulses on DAR sensitivity, *F*(2.32,116.13) = 68.69, *p* < .001, η_p_ ^2^ = .58; however, there was not a significant interaction between the number of pre-pulses and chronic drug pretreatment on DAR sensitivity, *F*(4.65,116.13) = 0.13, *p* = .98, η_p_ ^2^ = .01.

#### Baseline Dopamine Release and Half-life

Baseline dopamine release (µM) and synaptic half-life (sec) were compared between the three chronic drug pretreatment conditions before administration of the in-test drug challenge. With a representative release response in Figure 5A and means and standard error in Figure 5B, there was a significant main effect of chronic drug pretreatment on baseline dopamine release, *F*(2,51) = 3.21, *p* = .046, η^2^ = .11. Tukey’s post-hoc test revealed that mice chronically treated with ACEA had decreased dopamine release (*M* = 0.18, *SE* = 0.03) compared to mice chronically treated with vehicle (*M* = 0.29, *SE* = 0.03), *p* = .04. As can be seen in Figure 5C, baseline dopamine half-life was analyzed across the three drug pretreatment conditions. There was not a main effect of chronic drug pretreatment on baseline dopamine half-life, *F*(2,51) = 0.66, *p* = .52, η^2^ = .02.

**Figure 5.**
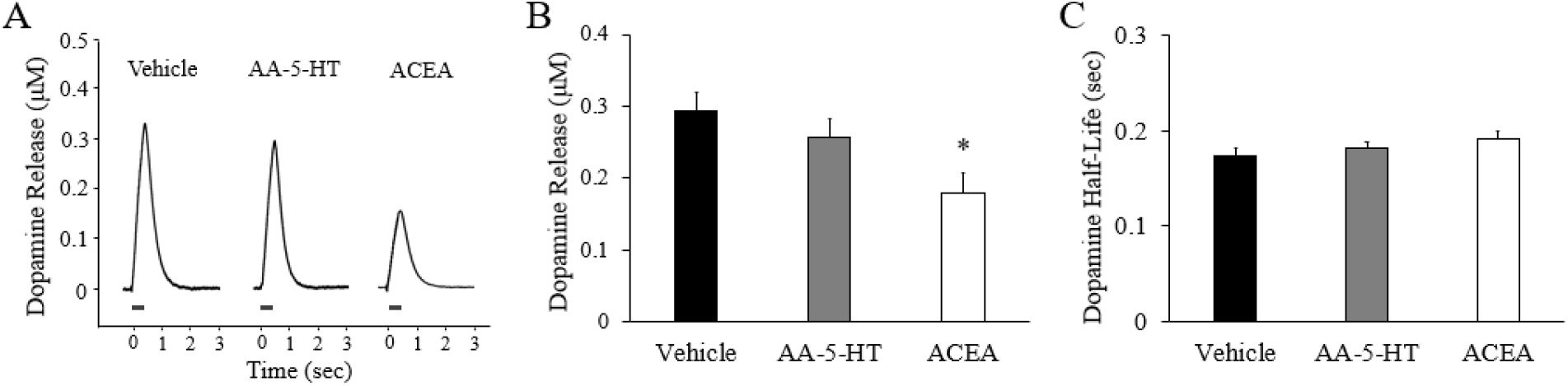
Baseline Dopamine Release and Half-Life. *Note.* (A) Representative stimulation-evoked dopamine release responses following chronic pretreatment with vehicle, arachidonoyl serotonin (AA-5-HT), or arachidonyl-2’-chloroethylamide (ACEA). (B) Dopamine release and (C) dopamine synaptic half-life in means ± SEMs following drug pretreatment. * indicates a significant decrease relative to vehicle pretreatment at *p* < .05.

#### Dopamine recordings following drug challenge (AA-5-HT)

During dopamine recordings, a subset of the mice chronically treated with AA-5-HT or vehicle received an i.p. drug challenge of either AA-5-HT (*n* = 7 and 6, respectively per chronic, drug treatment) or vehicle (*n* = 5 and 4, respectively per chronic, drug treatment). Percent change of dopamine release and half-life (with baseline dopamine release and half-life being 100%) were analyzed at 10 min intervals for 90 min post injection. As can be seen in Figure 6A, there was not a time point x drug pretreatment x drug challenge interaction effect, a time point x drug pretreatment interaction effect, a time point x drug challenge interaction effect, a main effect of chronic drug treatment, or a main effect of drug challenge on percent change in dopamine release (all *p*’s > .05). As can be seen in Figure 6B, there were also no interaction effects or main effects on percent change in dopamine half-life (all *p*’s > .05).

**Figure 6.**
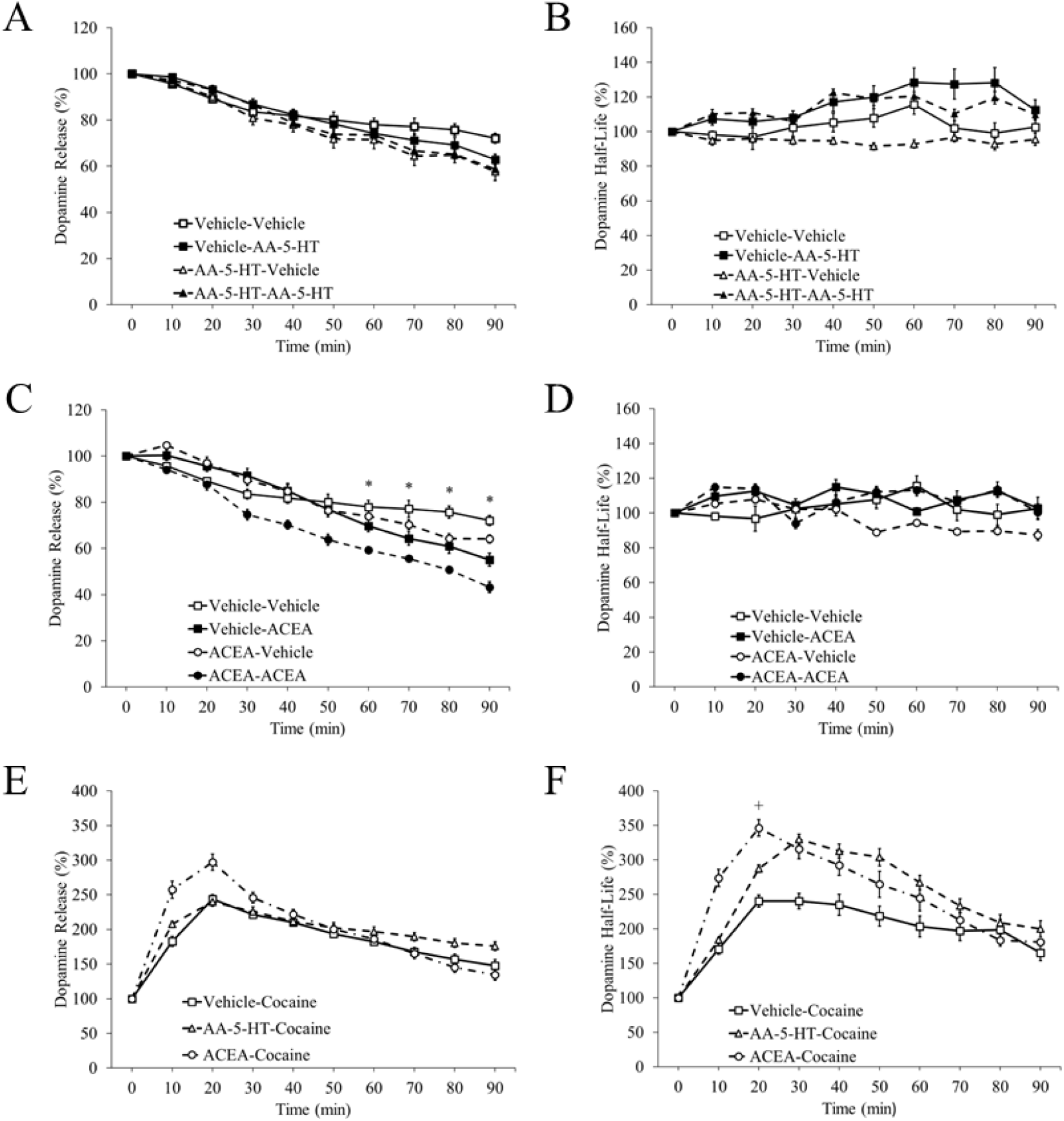
Drug Challenge Dopamine Release and Half-Life. *Note.* Percent change in dopamine release and half-life following drug challenges. The figure legend indicates the chronic, drug pretreatment then the drug challenge such that vehicle-vehicle represents mice that received vehicle pretreatment and vehicle drug challenge. Pretreatment of vehicle or arachidonoyl serotonin (AA-5-HT) on percent change in dopamine (A) release or (B) half-life presented as means ± SEMs. Pretreatment of vehicle or arachidonyl-2’-chloroethylamide (ACEA) on percent change in dopamine (C) release or (D) half-life presented as means ± SEMs. Percent change in dopamine (E) release and (F) half-life following cocaine administration in mice pretreated with vehicle, AA-5-HT, or ACEA. * indicates a significant difference between ACEA and vehicle challenge drug at *p* < .05 and + indicates a significant difference between ACEA and vehicle chronic, drug pretreatment at *p* < .05.

#### Dopamine Recordings following Drug Challenge (ACEA)

During dopamine recordings, a subset of the mice chronically treated with ACEA or vehicle received an i.p. drug challenge of either ACEA (*n* = 5 and 4, respectively per chronic, drug treatment) or vehicle (*n* = 5 and 4, respectively per chronic, drug treatment). Percent change of dopamine release and half-life (with baseline dopamine release and half-life being 100%) were analyzed at 10 min intervals for 90 min post injection. According to Mauchly’s test, the assumption of sphericity had been violated for time point, χ2(44) = 115.97, p < .001, and, therefore, degrees of freedom were corrected using Greenhouse-Geisser estimates of sphericity (ε = 0.43). As can be seen in Figure 6C, there was a time point x drug challenge interaction effect on percent change in dopamine release, *F*(3.89,54.49) = 4.70, *p* = .003, η_p_ ^2^ = .25. ACEA significantly reduced dopamine release in comparison to vehicle at 60, 70, 80, and 90 min post injection (all *p*’s < .05). However, there was no other interactions with time point or drug pretreatment or a main effect of drug pretreatment (all *p*’s > .05), indicating that the drug pretreatment did not influence the effect of the in-test ACEA injection. As can be seen in Figure 6D, there was not a time point x drug pretreatment x drug challenge interaction effect, a time point x drug pretreatment interaction effect, a time point x drug challenge interaction effect, a drug pretreatment x drug challenge interaction effect, a main effect of drug pretreatment, or a main effect of drug challenge on percent change in dopamine half-life (all *p*’s > .05).

#### Dopamine Recordings following Drug Challenge (Cocaine)

During dopamine recordings, a subset of the mice chronically treated with AA-5-HT, ACEA, or vehicle received an i.p. drug challenge of cocaine (*n* = 6, 5, and 6 per chronic, drug treatment group, respectively). Percent change of dopamine release and half-life (with baseline dopamine release being 100%) were analyzed at 10 min intervals for 90 min post injection. As can be seen in Figure 6E, there was not a time point x drug pretreatment interaction effect or main effect of drug pretreatment on percent change in dopamine release (both *p*’s > .05). As can be seen in Figure 6F, there was not a time point x drug pretreatment interaction effect or main effect of drug pretreatment on percent change in dopamine half-life (both *p*’s > .05). These data indicate that the chronic, drug pretreatments did not alter the timing of the dopaminergic response to cocaine. To assess differences at the peak effect time of cocaine, a one-way ANOVA was used to determine the effect of the chronic drug pretreatments on percent change in dopamine release and half-life at 20 min post-injection. No differences in percent change in dopamine release were observed between chronic, drug pretreatment groups (*p* > .05), but there was a significant main effect of drug pretreatment on percent change in dopamine half-life, *F*(2,14) = 6.39, *p* = .011, η_p_ ^2^ = .48. Tukey’s post-hoc test revealed that mice chronically pretreated with AA-5-HT did not differ from those pretreated with vehicle (*p* > .05); however, mice pretreated with ACEA had a greater percent change in dopamine half-life at 20 min post cocaine (*M* = 346.19%, *SE* = 26.77) compared to mice chronically pretreated with vehicle (*M* = 240.04%, *SE* = 21.68), *p* = .008.

### Experiment 2 (CPP and Saccharin Preference)

#### CPP

Time spent on the drug paired side and entries into each side were assessed for a bias on day one and for a difference between day eight (post-treatment) and day one (pretreatment). On the first day, the mice did not spend significantly more time on the drug-paired side (*M* = 911.08 s, *SE* = 31.91) than what was to be expected if the chamber was unbiased (i.e., 900 s for each side), *t*(26) = 1.62, *p* = .12. Time spent on the drug-paired side for day one was subtracted from time spent on the drug-paired side for day eight, and, as seen in Figure 7A, there was not a main effect of drug treatment on the difference, *F*(2,26) = 2.51, *p* = .10, η^2^ = .17. Similarly, the number of entries into the drug-paired side for day one was subtracted from the number of entries into the drug-paired side for day eight, and, as seen in Figure 7B, there was not a main effect of drug treatment on the difference, *F*(2,26) = 0.06, *p* = .94, η^2^ = .00.

**Figure 7.**
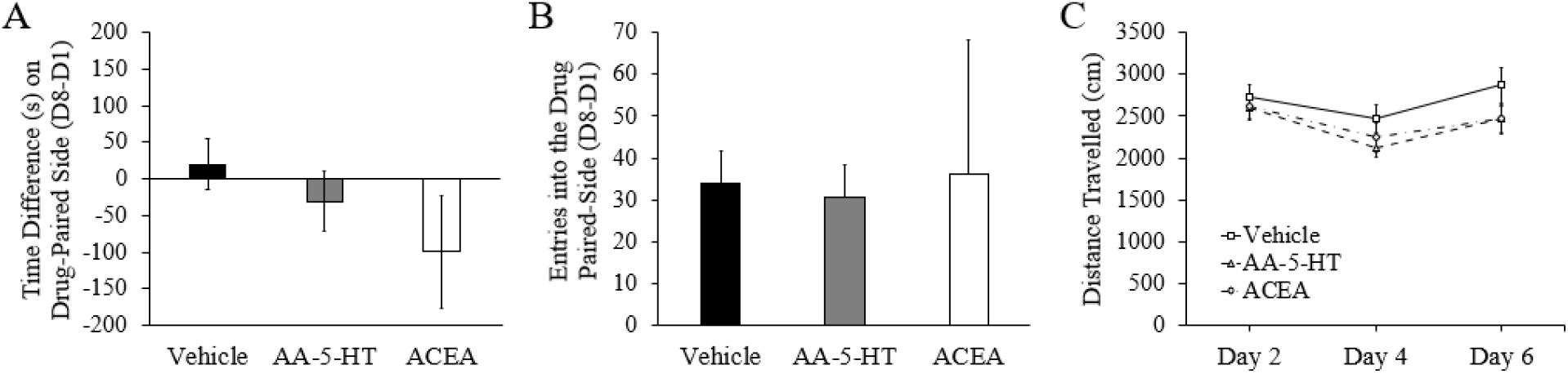
CPP. *Note.* Conditioned place preference data. (A) Drug challenge on time spent in the drug-paired side presented as day one (CPP pre-test) subtracted from day eight (CPP post-test). (B) Drug challenge on entries into the drug-paired side presented as day one (CPP pre-test) subtracted from day eight (CPP post-test). (C) Drug challenge on distance travelled on drug-paired days. CPP = conditioned place preference, AA-5-HT = arachidonoyl serotonin, and ACEA = arachidonyl-2’-chloroethylamide.

Distance travelled was recorded on the conditioning days (day two, four, and six) when mice were restricted to the drug-paired side of the apparatus and given either an i.p. injection of AA-5-HT, ACEA, or vehicle. As can be seen in Figure 7C, there was not a day x drug treatment interaction effect or main effect of drug treatment on distance travelled on drug-paired days (both *p*’s > .05).

#### Two-Bottle Choice Test

Mice were assessed for saccharin preference as a measure of anhedonia. As can be seen in Figure 8A, there was not a day x drug treatment interaction effect or main effect of drug treatment on saccharin consumed (both *p*’s > .05). Additionally, as can be seen in Figure 8B, there was not a day x drug treatment interaction effect or main effect of drug treatment on saccharin preference (both *p*’s > .05).

**Figure 8.**
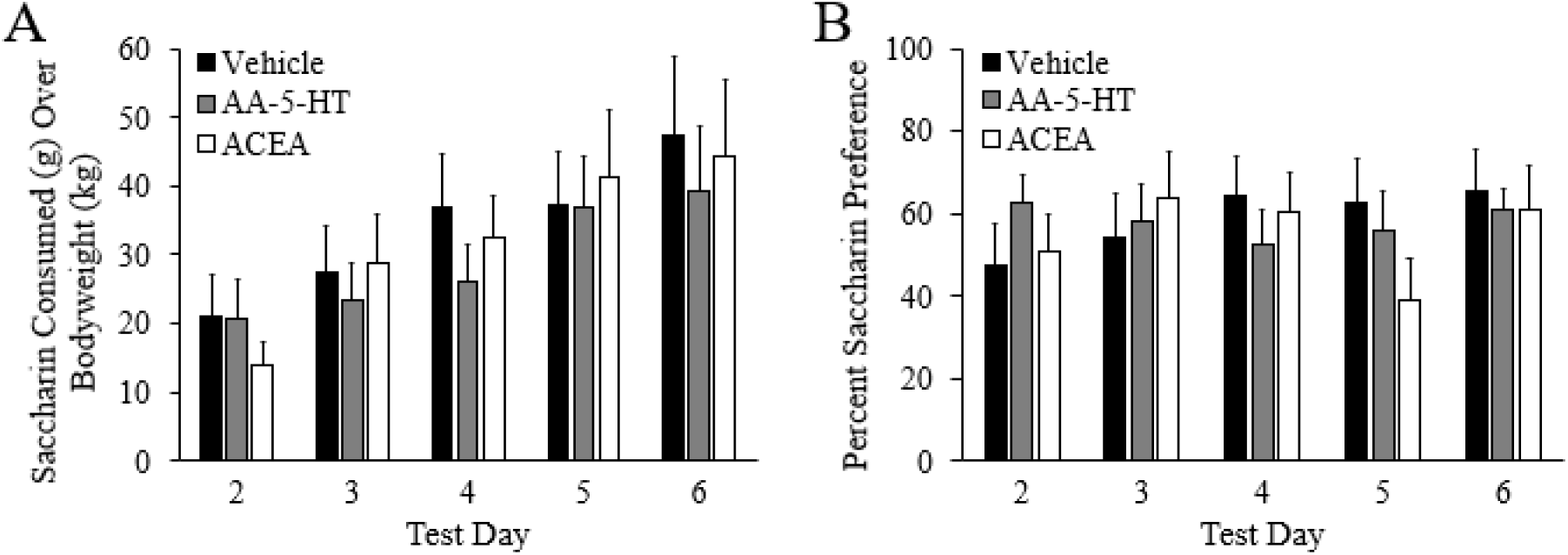
Saccharin Preference. *Note.* Two bottle saccharin preference data. (A) Drug challenge on saccharin consumed over body weight. (B) Drug challenge on saccharin consumed as a percent of total liquid consumed. AA-5-HT = arachidonoyl serotonin and ACEA = arachidonyl-2’-chloroethylamide.

## Discussion

The cannabinergic system is being researched as a pharmaceutical target for the treatment of several psychological disorders including anxiety, depression, and addiction (Batista et al., 2014; Bortolato et al., 2007; Chen et al., 1990; Cohen et al., 2002; de Mello Schier et al., 2014; Economidou et al., 2006; Fattore et al., 2005; Fogaça, et al., 2012; Freels et al., 2020; Gobbi et al., 2005; Hakimizadeh et al., 2012; Li et al., 2009; Patel & Hillard, 2006; Rey et al., 2012; Xi et al., 2008; Zaitone et al., 2012). The current study investigated the effects of chronic exposure to the indirect CB agonist AA-5-HT and the direct CB_1_R agonist ACEA on mesolimbic dopamine release and related behaviors. In Experiment 1, mice were pretreated with AA-5-HT, ACEA, or vehicle prior to assessments of exploratory behavior and dopamine functioning. In Experiment 2, mice underwent AA-5-HT, ACEA, or vehicle CPP then saccharin preference testing.

### Exploratory Behaviors

OF testing consisted of a baseline recording period, a drug injection (AA-5-HT, ACEA, or vehicle), and a post-drug recording period. Both human and animal models of substance use disorder indicate that novelty-seeking predicts the initiation of drug use and the transition to compulsive drug use (see Wingo et al., 2016). The chronic, drug pretreatments did not alter the baseline distance travelled, number of rears, or percent time in the center of the chamber, indicating that chronic exposure to the indirect or direct CB agonists did not influence the way in which the mice responded to the novel environment. The in-test AA-5-HT administration did not significantly alter OF measurements. Drug-induced locomotor activity is classically considered to be a predictor of whether the drug will be reinforcing (Wise & Bozarth, 1987); thus, the lack of change in distance travelled and rearing may be indicative of a non-addictive profile for AA-5-HT. Similarly, previous research has shown that neither AA-5-HT nor AACOCF_3_ (an FAAH inhibitor) altered locomotor activity in the OF (Freels et al., 2020; Rutkowska et al., 2006).

However, with no change in percent time spent in the center of the chamber, AA-5-HT also did not appear to have an anxiolytic effect. This may have occurred due to the choice of strain (B6) over a more anxious strain and/or due to the choice of test for anxiety, as moderate anxiolytic effects were found in BALB/c mice and in the elevated plus maze (Freels et al., 2020; Micale et al., 2009). Thus, more research is needed to explore potential strain and stress-context effects related to AA-5-HT administration.

An in-test injection of the direct CB_1_R agonist ACEA significantly decreased the rearing frequency of mice but did not alter distance travelled or percent time in center. Previous research has shown ACEA to not alter locomotor activities in the OF (Freels et al., 2020; Spiller et al., 2019). However, this may not be extendable to other CB_1_R agonists, as CB_1_R agonists have been shown to suppress both rearing and ambulation (Freels et al., 2020; Järbe et al., 2002; Rutkowska et al., 2006). The time length of the OF test may have affected the results in the current study. In general, mice moved infrequently between 70 and 90 min in the OF, and this time-induced floor effect may have masked drug-effects. Although more research is needed, the results of the current study suggest that these indirect (AA-5-HT) and direct (ACEA) CB agonists do not alter exploratory or anxiety-related behaviors as measured with OF.

### Dopamine Recordings

Amperometric measurements of dopamine consisted of a baseline recording period, drug injection (AA-5-HT, ACEA, or vehicle), and post-drug recording period. Chronic pretreatment with AA-5-HT did not significantly alter DAR sensitivity or dopamine dynamics (release or synaptic half-life) during the baseline recording period. The in-test AA-5-HT drug challenge did not significantly alter dopamine dynamics in either the saline or AA-5-HT pretreated groups.

These results support a lack of reinforcing effects of AA-5-HT administration via not increasing dopamine in the NAc, which coincides with previous studies exploring FAAH inhibition (Freels et al., 2020; Justinová et al., 2015). In the current study, chronic AA-5-HT pretreatment did not alter the dopaminergic response to cocaine; however, Mereu et al. (2013) found that pretreatment with the FAAH inhibitor URB597 facilitated cocaine-induced sensitization and increased percent change in dopamine levels in the NAc core. The current study differs from that of Mereu et al. (2013) in that mice were pretreated with chronic administration of AA-5-HT (rather than a single injection of URB597) and were only exposed to one injection of cocaine. These experimental differences and/or the dual FAAH inhibition and TRPV_1_ blocking action of AA-5-HT (vs. only FAAH inhibition by URB597) may have attenuated the alterations in dopamine signaling previously shown during a cocaine drug challenge in the NAc of mice. Altogether, the current results indicate that AA-5-HT treatment did not elicit a dopaminergic profile associated with abuse and addiction.

Mice chronically pretreated with ACEA, the direct CB_1_R agonist, did not differ from controls in DAR sensitivity or the synaptic half-life of dopamine but did display reduced dopamine release during baseline recordings (prior to the in-test drug challenges). The percent change in dopamine release was significantly decreased following the in-test ACEA injection compared to the vehicle injection in both chronically treated groups (vehicle and ACEA). These results mirror those from a previous study in our lab (Freels et al., 2020) and other studies utilizing FSCV to measure dopamine efflux following administration of direct CB_1_R agonists (Cheer et al., 2004; O’Neill et al., 2009). These findings suggest that ACEA does not elicit a dopaminergic profile associated with drug reinforcement and addiction; however, chronic pretreatment of ACEA did alter the dopaminergic response to cocaine. Mice that were pretreated with ACEA displayed an increased percent change in the synaptic half-life of dopamine following the in-test cocaine injection. Such findings support studies that have shown CB_1_R agonists increase the behavioral effects of psychostimulants (Gorriti et al., 1999; Lamarque et al., 2001) and have shown CB_1_R antagonists or inverse agonists decrease cocaine-induced changes in dopamine signaling (Mereu et al., 2013; Cheer et al., 2007; Wang et al., 2015). Altogether, the current results from amperometric recordings suggest that direct CB_1_R agonism by ACEA decreases dopamine release while potentially priming the mesolimbic dopamine system to have a respond more dramatically DAT inhibition.

### Conditioned Place Preference

During CPP, mice did not develop a CPP or CPA for AA-5-HT. Previous research exploring the CPP for drugs that prevent the breakdown of AEA found that AM404 (an AEA transporter inhibitor) and SBF126 (an inhibitor of fatty acid-binding protein 5 and fatty acid-binding protein 7) did not induce CPP in Wistar rats, Sprague-Dawley rats, or B6 mice (Bortolato et al., 2006; Scherma et al., 2012; Thanos et al., 2016).

Additionally, during CPP, mice did not develop a CPP or CPA for ACEA. Direct CB_1_R agonist administration has drug- and dose-dependent effects where, for example, THC has no effect at very low doses (around 1.0 mg/kg i.p. in mice.), a CPP at moderate doses (1.0 mg/kg i.p. in mice), and a CPA at high doses (starting at 5.0 mg/kg in mice) (Murray & Bevins, 2010; Panagis et al., 2014). Therefore, indirectly and directly agonizing the cannabinergic system can lead to CPP, but the doses in the current study, which were chosen due to their anxiolytic potential, did not, which suggests these doses are not rewarding or aversive in the current paradigm.

### Saccharin Preference Test

During the two-bottle choice test, mice treated with either AA-5-HT or ACEA did not develop a preference or aversion for saccharin. CP55940, URB597, and rimonabant had no effects on saccharin preference alone, but chronic, mild stress induced reductions in sucrose preference were attenuated by CP55940 or URB597 and enhanced by rimonabant (Bortolato et al., 2007; Rademacher & Hillard, 2007). This suggests that indirectly and directly agonizing the cannabinoid system does not alter preference for sweet rewards alone but can modify stress induced alterations in preference. Therefore, exploring preference for saccharin following both cannabinoid administration and stress may further the potential use of these drugs as anxiolytics. However, CP55940 increased progressive ratio responding for Ensure where rimonabant reduced it, suggesting the cannabinergic system still plays a role in the wanting of sweet rewards (Ward & Dykstra, 2005). AA-5-HT and ACEA administration did not alter saccharin preference, indicating these drugs do not elicit reward- or anhedonia-related behaviors.

### Cross-Modulation of the Cannabinergic and Dopaminergic Systems

Cross-modulation between neurotransmission systems can impact the efficacy and off-target effects of posited drugs. The current study explored the interplay between the cannabinergic system and dopaminergic system through the administration of AA-5-HT and ACEA. Through this study’s paradigms, chronic pretreatment with AA-5-HT did not alter behaviors related to dopamine in the OF, mirroring findings from our previous study with acute AA-5-HT administration (Freels et al., 2020). Also, AA-5-HT did not alter the measured aspects of phasic dopamine transmission (release, synaptic half-life, DAR sensitivity, and response to cocaine). Given that most drugs of abuse either directly or indirectly increase accumbal dopamine release, these findings indicate that AA-5-HT does not use the same neural mechanisms as would be expected for reinforcing drugs. Another FAAH inhibitor URB597 has been shown to increase cocaine-induced hyperlocomotion and cocaine-induced dopamine release in the NAc (Mereu et al., 2013), and AM404, a potent TRPV_1_ agonist and low affinity CB_1_R agonist, has been shown to reduce cocaine-induced hyperlocomotion (Vlachou et al., 2008). The dual actions of AA-5-HT (FAAH inhibition and TRPV_1_ agonism) may offset the effects on cocaine, as chronic pretreatment with AA-5-HT did not alter the dopaminergic effect of cocaine in the current study.

Cross-modulation between the cannabinergic and dopaminergic systems may be more greatly influenced by administration of drugs that directly bind to CB_1_Rs. In the current study, we did not find ACEA to alter distance travelled or rearing in the OF, and previous research from a laboratory found the same (Freels et al., 2020). However, we found that both chronic and acute injections of ACEA attenuate phasic dopamine release in the NAc, as has been similarly observed with another direct CB_1_R agonist (WIN55,212-2) in a study with FSCV (Cheer et al., 2004). Thus, there exists a cross-modulation between the cannabinergic and dopaminergic system in which agonism of the cannabinergic system attenuates mesolimbic dopamine release in the NAc, highlighting the importance of exploring the possible confounds of multi-drug treatments when exploring the efficacy of cannabinergic drugs. In the current study, chronic ACEA pretreatment led to an increased dopaminergic response to cocaine 20 min post-injection, adding to research that suggests the cannabinergic system may potentiate actions of psychostimulants (Gorriti et al., 1999; Lamarque et al., 2001; Mereu et al., 2013; Cheer et al., 2007; Wang et al., 2015). However, it should be noted that CB_1_R agonists have also been shown to have the opposite effect, reducing psychostimulant-induced behaviors (Vlachou et al., 2003; Panlilio et al., 2007). As such, more research is needed to understand the cross-modulation between the cannabinoid and dopamine systems. The current study is limited by the use of only adult, male mice. The effects of cannabinergic drugs on dopamine functioning may be dependent on both age and sex. For example, previous behavioral studies have shown that cannabinoid agonists administered during adolescence enhance the reinforcing effects of cocaine to a greater degree than those administered in adulthood (Dow-Edwards & Izenwasser, 2012, Friedman et al., 2019). Although sex-dependent effects of the rewarding effects of cannabinoids are under researched and often contradictory (see Calakos et al., 2017), female cannabis users have been shown to progress from first use to cannabis use disorder more rapidly than males (Hernandez-Avila et al., 2004; Kahn et al., 2013), and female rats have been shown to display faster acquisition of cannabinoid self-administration than males (Scherma et al., 2016).

## Conclusions

As the cannabinergic system remains a target for new pharmaceuticals, further studies on the influence of cannabinergic drugs on neural systems underlying reward and motivation is crucial. The current findings suggest that indirect agonism of the cannabinoid system (through combined FAAH inhibition and TRPV_1_ antagonism by AA-5-HT) has less of an effect on mesolimbic dopamine release and related behaviors than direct CB_1_R agonism (by ACEA). Neither drug (AA-5-HT or ACEA) directly increased dopamine release or induced behaviors associated with drug reinforcement (drug-induced locomotor activity or CPP). Instead, ACEA decreased dopamine release but assumedly not to an extent that would induce anhedonia given that saccharin preference was not altered. Thus, neither drug induced behavioral or neurochemical indications of reinforcement that would lead to abuse liability. Pretreatment with ACEA did increase the dopaminergic effect of cocaine 20 min post-injection while pretreatment with AA-5-HT did not. Altogether, our results suggest that direct CB_1_R agonism may play more of a role in mediating reward and motivation than indirect CB agonism through FAAH inhibition and TRPV_1_ antagonism. More research is needed to generalize findings on the interactions between cannabinergic and dopaminergic systems across drug mechanisms and population characteristics (such as age and sex).

## Notes

### Competing Interest Statement

The authors have declared no competing interest.

### Summary of Updates

Figures were combined to be more concise, and the text was edited for clarity and an improved explanation of the findings.

## References

Batista, L.A., Gobira, P.H., Viana, T.G., Aguiar, D.C., & Moreira, F.A. (2014). Inhibition of endocannabinoid neuronal uptake and hydrolysis as strategies for developing anxiolytic drugs. Behavioural Pharmacology, 25(5-6), 425–433. https://doi.org/10.1097/FBP.0000000000000073

Berridge, K.C. & Robinson, T.E. (1998). What is the role of dopamine in reward: Hedonic impact, reward learning, or incentive salience? Brain Research Reviews, 28(3), 309–369. https://doi.org/10.1016/S0165-0173(98)00019-8

Blankman, J.L. & Cravatt, B.F. (2013). Chemical probes of endocannabinoid metabolism. Pharmacological Reviews, 65(2), 849–871. https://doi.org/10.1124/pr.112.006387

Blum, K., Khalsa, J., Cadet, J.L., Baron, D., Bowirrat, A., Boyett, B, Lott, L., Brewer, R., Gondré-Lewis, M., Bunt, G., Kazmi, S., & Gold, M.S. (2021). Cannabis-induced hypodopaminergic anhedonia and cognitive decline in humans: Embracing putative induction of dopamine homeostasis. Frontiers in Psychiatry, 12, 623403. https://doi.org/10.3389/fpsyt.2021.623403

Bortolato, M., Campalongo, P., Mangieri, R.A., Scattoni, M.L., Frau, R., Trezza, V., La Rana, G., Russo, R., Calignano, A., Gessa, G.L., Cuomo, V., & Piomelli, D. (2006). Anxiolytic-like properties of anandamide transport inhibitor AM404. Neuropsychopharmacology, 31(12), 2652–2659. https://doi.org/10.1038/sj.npp.1301061

Bortolato, M., Mangieri, R.A., Fu, J., Kim, J.H., Arguello, O., Duranti, A., Tontini, A., Mor, M., Tarzia, G., & Piomelli, D. (2007). Antidepressant-like activity of the fatty acid amide hydrolase inhibitor URB597 in a rat model of chronic mild stress. Biological Psychiatry, 62(10), 1103–1110. https://doi.org/10.1016/j.biopsych.2006.12.001

Cheer, J.F., Wassum, K.M., Heien, M.L., Phillips, P.E., & Wightman, R.M. (2004). Cannabinoids enhance subsecond dopamine release in the nucleus accumbens of awake rats. The Journal of Neuroscience, 24(18), 4393–4400. https://doi.org/10.1523/JNEUROSCI.0529-04.2004

Cheer, J.F., Wassum, K.M., Sombers, L.A., Heien, M.L., Ariansen, J.L., Aragona, B.J., Pillips, P.E., & Wightman, R.M. (2007). Phasic dopamine release evoked by abused substances requires cannabinoid receptor activation. The Journal of Neuroscience, 27(4), 791–795. https://doi.org/10.1523/JNEUROSCI.4152-06.2007

Chen, J.P. Paredes, W., Li, Y., Smith, D., Lowinson, J., & Gardner, E.L. (1990). Delta 9-tetrahydrocannabinol produces naloxone-blockable enhancement of presynaptic basal dopamine efflux in the nucleus accumbens of conscious, freely-moving rats as measured by intracerebral microdialysis. Psychopharmacology, 102(2). https://doi.org/10.1007/BF02245916.

Cohen, C., Perrault, G., Voltz, C., Steinberg, R., & Soubrié, P. (2002). SR141716, a central cannabinoid (CB_1_) receptor antagonist, blocks the motivational and dopamine-releasing effects of nicotine in rats. Behavioural Pharmacology, 13(5-6), 451–463. https://doi.org/10.1097/00008877-200209000-00018

Day, J.J., Roitman, M.F., Wightman, R.M., & Carelli, R.M. (2007). Associative learning mediates dynamic shifts in dopamine signaling in the nucleus accumbens. Nature Neuroscience, 10(8), 1020–1028. https://doi.org/10.1038/nn1923

de Mello Schier, A.R., de Oliveira Ribeiro, N.P., Coutinho, D.S., Machado, S., Arias-Carrión, O., Crippa, J.A., Zuardi, A.W., Nardi, A.E., & Silva, A.C. (2014). Antidepressant-like and anxiolytic-like effects of cannabidiol: A chemical compound of Cannabis sativa. CNS & Neurological Disorders − Drug Targets, 13(6), 953–60. https://doi.org/10.2174/1871527313666140612114838

De Petrocellis, L., Bisogno, T., Maccarrone, M., Davis, J.B., Finazzi-Agro, A., & Di Marzo, V. (2001). The activity of anandamide at vanilloid VR1 receptors requires facilitated transport across the cell membrane and is limited by intracellular metabolism. Journal of Biological Chemistry, 276(16), 12856–12863. https://doi.org/10.1074/jbc.M008555200

De Vries, T. J., Shaham, Y., Homberg, J. R., Crombag, H., Schuurman, K., Dieben, J., Vanderschuren, L. J., & Schoffelmeer, A. N. (2001). A cannabinoid mechanism in relapse to cocaine seeking. Nature Medicine, 7(10), 1151–1154. https://doi.org/10.1038/nm1001-1151

Dinh, T.P., Carpenter, D., Leslie, F.M., Freund, T.F., Katona, I., Sensi, S.L., Kathuria, S., & Piomelli, D. (2002). Brain monoglyceride lipase participating in endocannabinoid inactivation. Proceedings of the National Academy of Sciences of the United States of America, 99(16), 10819–10824. https://doi.org/10.1073/pnas.152334899

Dow-Edwards, D., & Izenwasser, S. (2012). Pretreatment with Δ9-tetrahydrocannabinol (THC) increases cocaine-stimulated activity in adolescent but not adult male rats. Pharmacology Biochemistry and Behavior, 100(3), 587–591. https://doi.org/10.1016/j.pbb.2011.09.003

Dugast, C., Suaud-Chagny, M.F., & Gonon, F. (1994). Continuous in vivo monitoring of evoked dopamine release in the rat nucleus accumbens by amperometry. Neuroscience, 62, 647–654. https://doi.org/10.1016/0306-4522(94)90466-9

Economidou, D., Mattioli, L., Cifani, C., Perfumi, M., Massi, M., Cuomo, V., Trabace, L., & Ciccocioppo, R. (2006). Effect of the cannabinoid CB_1_ receptor antagonist SR-141716A on ethanol self-administration and ethanol-seeking behaviour in rats. Psychopharmacology, 183(4), 394–403. https://doi.org/10.1007/s00213-005-0199-9

Everitt, B.J. & Robbins, T.W. (2005). Neural systems of reinforcement for drug addiction: From actions, to habits, to compulsion. Nature Neuroscience, 8, 1481–1489. https://doi.org/10.1038/nn1579

Fadda, P., Scherma, M., Spano, M.S., Salis, P., Melis, V, Fattore, L., & Fratta, W. (2006). Cannabinoid self-administration increases dopamine release in the nucleus accumbens. NeuroReport, 17(15), 1629–1632. https://doi.org/10.1097/01.wnr.0000236853.40221.8e

Fattore, L., Martellotta, M. C., Cossu, G., Mascia, M. S., & Fratta, W. (1999). CB_1_ cannabinoid receptor agonist WIN 55,212-2 decreases intravenous cocaine self-administration in rats. Behavioural Brain Research, 104(1-2), 141–146. https://doi.org/10.1016/s0166-4328(99)00059-5

Fattore, L., Spano, S., Cossu, G., Deiana, S., Fadda, P., & Fratta, W. (2005). Cannabinoid CB_1_ antagonist SR 141716A attenuates reinstatement of heroin self-administration in heroin-abstinent rats. Neuropharmacology, 48(8), 1097–1104. https://doi.org/10.1016/j.neuropharm.2005.01.022

Fielding, J.R., Rogers, T.D., Meyers, A.E., Miller, M.M., Nelms, J.L., Mittleman, G., Blaha, C.D., & Sable, H.J. (2013). Stimulation-evoked dopamine release in the nucleus accumbens following cocaine administration in rats perinatally exposed to polychlorinated biphenyls. Toxicology Sciences, 136(1), 144–153. https://doi.org/10.1093/toxsci/kft171

Fogaça, M.V., Aguiar, D.C., Moreira, F.A., Guimarães, F.S. (2012). The endocannabinoid and endovanilloid systems interact in the rat prelimbic medial prefrontal cortex to control anxiety-like behavior. Neuropharmacology, 63(2), 202–210. https://doi.org/10.1016/j.neuropharm.2012.03.007

Freels, T.F., Lester, D.B., & Cook, M.N. (2019). Arachidonoyl serotonin (AA-5-HT) modulates general fear-like behavior and inhibits mesolimbic dopamine release. Behavioural Brain Research, 362, 140–151. https://doi.org/10.1016/j.bbr.2019.01.010

Friedman, A. L., Meurice, C., & Jutkiewicz, E. M. (2019). Effects of adolescent Δ9-tetrahydrocannabinol exposure on the behavioral effects of cocaine in adult Sprague-Dawley rats. Experimental and Clinical Psychopharmacology, 27(4), 326–337. https://doi.org/10.1037/pha0000276

Giang, D.K. & Cravatt, B.F. (1997). Molecular characterization of human and mouse fatty amide hydrolases. Proceedings of the National Academy of Sciences of the United States of America, 94(6), 2238–2242. https://doi.org/10.1073/pnas.94.6.2238

Gobbi, G., Bambico, F.R., Mangieri, R., Bortolato, M., Campolongo, P., Solinas, M., Cassano, T., Morgerse, M.G., Debonnel, G., Duranti, A., Tontini, A., Tarzia, G., Mor, M., Trezza, V., Goldberg, S.R., & Piomelli, D. (2005). Antidepressant-like activity and modulation of brain monoaminergic transmission by blockade of anandamide hydrolysis. Proceedings of the National Academy of Sciences of the United States of America, 102(51), 18620–18625. https://doi.org/10.1073/pnas.0509591102

Goto, Y., Otani, S., & Grace, A. A. (2007). The Yin and Yang of dopamine release: A new perspective. Neuropharmacology, 53(5), 583–587. https://doi.org/10.1016/j.neuropharm.2007.07.007

Grace, A. A., Floresco, S. B., Goto, Y., & Lodge, D. J. (2007). Regulation of firing of dopaminergic neurons and control of goal-directed behaviors. Trends in Neurosciences, 30(5), 220–227. https://doi.org/10.1016/j.tins.2007.03.003

Gulyas, A.I., Cravatt, B.F., Bracey, M.H., Dinh, T.P., Piomelli, D., Boscia, F., & Freund, T.F. (2004). Segregation of two endocannabinoid-hydrolyzing enzymes into pre- and postsynaptic compartments in the rat hippocampus, cerebellum, and amygdala. The European Journal of Neuroscience, 20(2), 441–458. https://doi.org/10.1111/j.1460-9568.2004.03428.x

Haj-Dahmane, S., Shen, R.-Y., Elmes, M.W., Studholme, K., Kanjiya, M.P., Bodgan, D., Thanos, P.K., Miyauchi, J.T., Tsirka, S.E., & Kaczocha, M. (2018). Fatty-acid-binding protein 5 controls retrograde cannabinoid signaling at central glutamate synapses. Proceedings of the National Academy of Sciences of the United States of America, 115(3), 3482–3487. https://doi.org/10.1073/pnas.1721339115

Hakimizadeh, E., Oryan, S., Moghaddam, A.H., Shamsizadeh, A., & Roohbakhsh, A. (2012). Endocannabinoid system and TRPV1 receptors in the dorsal hippocampus of the rats modulate anxiety-like behaviors. Iranian Journal of Basic Medical Sciences, 15(3), 795–802. https://doi.org/10.22038/ijbms.2012.4863

Han, X., He, Y., Bi, G.H., Zhang, H.Y., Song, R., Liu, Q.R., Egan, J.M., Gardner, E.L., Li, J., & Xi, Z.X. (2017). CB1 receptor activation on VgluT2-expressing glutamatergic neurons underlies Δ9-tetrahydrocannabinol (Δ9-THC)-induced aversive effects in mice. Scientific Reports, 7(1), 12315. https://doi.org/10.1038/s41598-017-12399-z

Holloway, Z., Freels, T.F., Comstock, J.F., Nolen, H.G., Sable, H.J. & Lester, D.B. (2018). Comparing phasic dopamine dynamics in the striatum, nucleus accumbens, amygdala, and medial prefrontal cortex. Synapse, 73(2). https://doi.org/10.1002/syn.22074

Howlett, A.C., Barth, F., Bonner, T.I., Cabral, G., Casellas, P., Devane, W.A., Felder, M., Herkenham, M., Mackie, K., Martin, B.R., Mechoulam, R., & Pertwee, R.G. (2002). International union of pharmacology. Xxvii. Classification of cannabinoid receptors. Pharmacological Reviews, 54(2), 161–202. https://doi.org/10.1124/pr.54.2.161

Hyland, B. I., Reynolds, J. N., Hay, J., Perk, C. G., & Miller, R. (2002). Firing modes of midbrain dopamine cells in the freely moving rat. Neuroscience, 114(2), 475–492. https://doi.org/10.1016/s0306-4522(02)00267-1

Järbe, T.U., Andrzejewski, M.E., & DiPatrizio, N.V. (2002). Interactions between the CB1 receptor agonist Delta 9-THC and the CB1 receptor antagonist SR-141716 in rats: Open-field revisited. Pharmacology, Biochemistry, and Behavior, 73(4), 911–919. https://doi.org/10.1016/S0091-3057(02)00938-3

Justinová, Z., Panlilio, L. V., Moreno-Sanz, G., Redhi, G. H., Auber, A., Secci, M. E., Mascia, P., Bandiera, T., Armirotti, A., Bertorelli, R., Chefer, S. I., Barnes, C., Yasar, S., Piomelli, D., & Goldberg, S. R. (2015). Effects of fatty acid amide hydrolase (FAAH) inhibitors in non-human primate models of nicotine reward and relapse. Neuropsychopharmacology, 40(9), 2185–2197. https://doi.org/10.1038/npp.2015.62

Kaczocha, M., Glaser, S.T., & Deutsch, D.G. (2009). Identification of intracellular carriers for the endocannabinoid anandamide. Proceedings of the National Academy of Sciences of the United States of America, 106(15), 6375–6380. https://doi.org/10.1073/pnas.0901515106

Kano, M., Ohno-Shosaku, T., Hashimotodani, Y., Uchigashima, M., & Watanabe, M. (2009). Endocannabinoid-mediated control of synaptic transmission. Physiological Review, 89(1), 309-380. https://doi.org/10.1152/physrev.00019.2008

Kaur, J.A. & Gibson, H.E. (2009). Hot flash: TRPV channels in the brain. Trends in Neurosciences, 32(4), 215–224. https://doi.org/10.1016/j.tins.2008.12.006

Lecca, D., Cacciapaglia, F., Valentini, V., & Di Chiara, G. (2006). Monitoring extracellular dopamine in the rat nucleus accumbens shell and core during acquisition and maintenance of intravenous WIN 55,212-2 self-administration. Psychopharmacology, 188(1), 63–74. https://doi.org/10.1007/s00213-006-0475-3

Li, X., Hoffman, A. F., Peng, X. Q., Lupica, C. R., Gardner, E. L., & Xi, Z. X. (2009). Attenuation of basal and cocaine-enhanced locomotion and nucleus accumbens dopamine in cannabinoid CB1-receptor-knockout mice. Psychopharmacology, 204(1), 1–11. https://doi.org/10.1007/s00213-008-1432-0

Lupica, C. R., & Riegel, A. C. (2005). Endocannabinoid release from midbrain dopamine neurons: A potential substrate for cannabinoid receptor antagonist treatment of addiction. Neuropharmacology, 48(8), 1105–1116. https://doi.org/10.1016/j.neuropharm.2005.03.016

Maccarrone, M. (2017). Metabolism of the endocannabinoid anandamide: Open questions after 25 years. Frontiers in Molecular Neuroscience, 10, 166, https://doi.org/10.3389/fnmol.2017.00166

Maione, S., De Petrocellis, L., de Novellis, V., Moriello, A.S., Petrosino, S., Palazzo, E., Rossi, F.S., & Woodward, D.F., & Di Marzo, V. (2007) Analgesic actions of *N-*arachidonoyl-serotonin, a fatty acid amide hydrolase inhibitor with antagonistic activity at vanilloid TRPV1 receptors. British Journal of Pharmacology, 150(6), 766–781. https://doi.org/10.1038/sj.bjp.0707145

Mereu, M., Tronci, V., Chun, L.E., Thomas, A.M., Green, J.L., Katz, J.L., & Tanda G. (2013). Cocaine-induced endocannabinoid release modulates behavioral and neurochemical sensitization in mice. Addiction Biology, 20(1), 91–103. https://doi.org/10.1111/adb.12080

Micale, V., Cristino, L. Tamburella, A., Petrosino, S., Leggio, G.M., Drago, F., & Di Marzo, V. (2009). Anxiolytic effects in mice of a dual blocker of fatty acid amide hydrolase and transient receptor potential type-1 channels. Neuropsychopharmacology, 34(3), 593–606. https://doi.org/10.1038/npp.2008.98

Mittleman, G., Call, S.B., Cockroft, J.L., Goldowitz, D., Matthews, D.B., & Blaha, C.D. (2011). Dopamine dynamics associated with, and resulting from, schedule-induced alcohol self-administration: Analyses in dopamine transporter knockout mice. Alcohol, 45(4), 325–339. https://doi.org/10.1016/j.alcohol.2010.12.006

Montague, P.R., Hyman, S.E., & Cohen, J.D. (2004). Computational roles for dopamine in behavioural control. Nature, 431, 760–767. https://doi.org/10.1038/nature03015

Murray, J.E. & Bevins, R.A. (2010). Cannabinoid conditioned reward and aversion: Behavioral and neural processes. ACS Chemical Neuroscience, 1(4), 265–278. https://doi.org/10.1021/cn100005p

National Institute of Health. (2012). The Public Health Service policy on humane care and use of laboratory animals. US Department of Health and Human Service.

National Research Council (2003). Guidelines for the care and use of mammals in neuroscience and behavioral research. The National Academies Press.

O’Neill, C., Evers-Donnelly, A., Nicholson, D., O’Boyle, K. M., & O’Connor, J. J. (2009). D_2_ receptor-mediated inhibition of dopamine release in the rat striatum in vitro is modulated by cb1 receptors: Studies using fast cyclic voltammetry. The Journal of Neurochemistry, 108(3), 545–551. https://doi.org/10.1111/j.1471-4159.2008.05782.x

Ohno-Shosaku, T., Maejima, T., & Kano, M. (2001). Endogenous cannabinoids mediate retrograde signals from depolarized postsynaptic neurons to presynaptic terminals. Neuron, 29(3), 729–738. https://doi.org/10.1016/S0896-6273(01)00247-1

Panagis, G., Mackey, B., & Vlachou, S. (2014). Cannabinoid regulation of brain reward processing with an emphasis on the role of CB_1_ receptors: A step back into the future. Frontiers in Psychiatry, 5, 92. https://doi.org/10.3389/fpsyt.2014.00092

Patel, S. & Hillard, C.J. (2006). Pharmacological evaluation of cannabinoid receptor ligands in a mouse model of anxiety: Further evidence of an anxiolytic role for endogenous cannabinoid signaling. The Journal of Pharmacology and Experimental Therapeutics, 318(1), 304–311. https://doi.org/10.1124/jpet.106.101287

Paxinos, G. & Franklin, K.B. (2001). The mouse brain in stereotaxic coordinates (2nd ed.). San Diego, CA: Academic Press.

Peters, K., Oleson, E., Covey, D., Mateo, Y., Tonini, R., & Cheer, J.F. (2021). Endocannabinoid modulation of the dopamine System: Implications for motivated behavior. Biological Psychiatry, 89(9), S52–S53. https://doi.org/10.1016/j.biopsych.2021.02.148

Prater, W.T., Swamy, M., Beane, M.D., & Lester, D.B. (2018). Examining the effects of common laboratory methods on the sensitivity of carbon fiber electrodes in amperometric recordings of dopamine. Journal of Behavioral and Brain Science, 8(3), 117–125. https://doi.org/10.4236/jbbs.2018.83007

Rademacher, D.J. & Hillard, C.J. (2007). Interactions between endocannabinoids and stress-induced decreased sensitivity to natural reward. Progress in Neuro-psychopharmacology & Biological Psychiatry, 31(3), 633–641. https://doi.org/10.1016/j.pnpbp.2006.12.013

Rey, A.A., Purrio, M., Viveros, M.P., & Lutz, B. (2012) Biphasic effects of cannabinoids in anxiety responses: CB1 and GABA_B_ receptors in the balance of GABAergic and glutamatergic neurotransmission. Neuropsychopharmacology, 37(12), 2624–234. https://doi.org/10.1038/npp.2012.123

Rosenbaum, T. & Simon, S.A. (2007). TRPV_1_ receptors and signal transduction. In, W.B. Liedtke & Heller S. (Eds.) TRP ion channel function in sensory transduction and cellular signaling cascades (pp. 69–85). Boca Raton, FL: CRC Press: Taylor and Francis Group.

Ross, R.A. (2003). Anandamide and vanilloid TRPV1 receptors. British Journal of Pharmacology, 140(5), 790–801. https://doi.org/10.1038/sj.bjp.0705467

Rutkowska, M., Jamontt, J., & Gliniak, H. (2006). Effects of cannabinoids on the anxiety-like response in mice. Pharmacological Reports, 58(2), 200–206.

Scherma, M., Justinová, Z., Zanettini, C., Panlilio, L.V., Mascia, P. Fadda, P., Fratta, W., Makriyannis, A., Vadivel, S.K., Gamaleddin, I., & Le Foll, B., & Goldberg, S.R. (2012). The anandamide transport inhibitor AM404 reduces the rewarding effects of nicotine and nicotine-induced dopamine elevations in the nucleus accumbens shell in rats. British Journal of Pharmacology, 165(8), 2539–2548. https://doi.org/10.1111/j.1476-5381.2011.01467.x

Schultz, W., Apicella, P., & Ljungberg, T. (1993). Responses of monkey dopamine neurons to reward and conditioned stimuli during successive steps of learning a delayed response task. The Journal of Neuroscience, 13(3), 900–913. https://doi.org/10.1523/JNEUROSCI.13-03-00900.1993

Sheffield, F.D. & Roby, T.B. (1950). Reward value of a non-nutritive sweet taste. Journal of Comparative and Physiological Psychology, 43(6), 471–481. https://doi.org/10.1037/h0061365

Smart, D., Gunthorpe, M.J., Jerman, J.C., Nasir, S., Gray, J., Muir, A.I., Chambers, J.K., Randall, A.D., & Davis, J.B. (2000). The endogenous lipid anandamide is a full agonist at the human vanilloid receptor (hVR1). British Journal of Pharmacology, 129(2), 227–230. https://doi.org/10.1038/sj.bjp.0703050

Solinas, M., Justinová, Z., Goldberg, S.R., & Tanda, G. (2006). Anandamide administration alone and after inhibition of fatty acid amide hydrolase (FAAH) increases dopamine in the nucleus accumbens shell in rats. Journal of Neurochemistry, 98(2), 408–419. https://doi.org/10.1111/j.1471-4159.2006.03880.x

Soria, G., Mendizábal, V., Touriño, C., Robledo, P., Ledent, C., Parmentier, M., Maldonado, R., & Valverde, O. (2005). Lack of CB1 cannabinoid receptor impairs cocaine self-administration. Neuropsychopharmacology, 30(9), 1670–1680. https://doi.org/10.1038/sj.npp.1300707

Spiller, K.J., Bi, G-h., He, Y., Galaj, E., Gardner, E.L., & Xi, Z.-X. (2019). Cannabinoid CB_1_ and CB_2_ receptor mechanisms underlie cannabis reward and aversion. British Journal of Pharmacology, 176(9), 1268–1281. https://doi.org/10.1111/bph.14625

Szabo, B., Siemes, S., & Wallmichrath, I. (2002). Inhibition of GABAergic neurotransmission in the ventral tegmental area by cannabinoids. The European Journal of Neuroscience, 15(12), 2057–2061. https://doi.org/10.1046/j.1460-9568.2002.02041.x

Tanda, G., Pontieri, F.E., & Di Chiara, G. (1997). Cananbinoid and heroin activation of mesolimbic dopamine transmission by a common μ_1_ opioid receptor mechanism. Science, 276(5321), 2048–2050. https://doi.org/10.1126/science.276.5321.2048

Thanos, P.K., Clavin, B.H., Hamilton, J., O’Rourke, J.R., Maher, T., Koumas, C., Miao, E., Lankop, J., Elhage, A., Haj-Dahmane, S., Deutsch, D., & Kaczocha, M. (2016). Examination of the addictive and behavioral properties of fatty acid-binding protein inhibitor SBFI26. Frontiers in Psychiatry, 7, 54. https://doi.org/10.3389/fpsyt.2016.00054

Vlachou, S., Nomikos, G. G., & Panagis, G. (2003). WIN 55,212-2 decreases the reinforcing actions of cocaine through CB_1_ cannabinoid receptor stimulation. Behavioural Brain Research, 141(2), 215–222. https://doi.org/10.1016/s0166-4328(02)00370-4

Vlachou, S., Stamatopoulou, F., Nomikos, G.G., & Panagis, G. (2008). Enhancement of endocannabinoid neurotransmission through CB_1_ cannabinoid receptors counteracts the reinforcing and psychostimulant effects of cocaine. International Journal of Neuropsychopharmacology, 11(7), 905–923. https://doi.org/10.1017/S1461145708008717

Wang, H., Treadway, T., Covey, D.P., Cheer, J.F., & Lupica, C.R. (2015). Cocaine-induced endocannabinoid mobilization in the ventral tegmental area. Cell Reports, 12(12), 1997–2008. https://doi.org/10.1016/j.celrep.2015.08.041

Ward, S.J. & Dykstra, L.A. (2005). The role of CB_1_ receptors in sweet versus fat reinforcement: Effect of CB_1_ receptor deletion, CB_1_ receptor antagonism (SR141716A) and CB_1_ receptor agonism (CP-55940). Behavioural Pharmacology, 16(5-6), 381–388. https://doi.org/10.1097/00008877-200509000-00010

Wingo, T., Nesil, T., Choi, J. S., & Li, M. D. (2016). Novelty seeking and drug addiction in humans and animals: From behavior to molecules. Journal of Neuroimmune Pharmacology, 11(3), 456–470. https://doi.org/10.1007/s11481-015-9636-7

Winters, B.D., Krüger, J.M., Huang, X., Gallaher, Z.R., Ishikawa, M., Czaja, K., Krueger, J.M., Huang, Y.H., Schlüter, O.M., & Dong, Y. (2012). Cannabinoid receptor 1-expressing neurons in the nucleus accumbens. Proceedings of the National Academy of Sciences of the United States of America, 109(40), 2717–2725. https://doi.org/10.1073/pnas.1206303109

Wise, R.A. & Bozarth, M.A. (1987) A psychomotor stimulant theory of addiction. Psychological Review, 94(9), 469–492.

Xi, Z. X., Spiller, K., Pak, A. C., Gilbert, J., Dillon, C., Li, X., Peng, X. Q., & Gardner, E. L. (2008). Cannabinoid CB1 receptor antagonists attenuate cocaine’s rewarding effects: experiments with self-administration and brain-stimulation reward in rats. Neuropsychopharmacology, 33(7), 1735–1745. https://doi.org/10.1038/sj.npp.1301552

Zaitone, S.A., El-Wakeil, A.F., & Abou-El-Ela, S.H. (2012). Inhibition of fatty acid amide hydrolase by URB597 attenuates the anxiolytic-like effect of acetaminophen in the mouse elevated plus-maze test. Behavioural Pharmacology, 23(4), 417–425. https://doi.org/10.1097/FBP.0b013e3283566065

